# Real-Time Viability Assessment of Ex Vivo Mouse Kidneys for Transplant Applications Using Dynamic Optical Coherence Tomography

**DOI:** 10.1101/2025.08.30.673280

**Authors:** Ke Zhang, Feng Yan, Trisha Valerio, Cui Yan, Qinghao Zhang, Ronghao Liu, Xiaoyu Ma, Junyuan Liu, Chen Wang, Tri Vu, Bornface Mutembei, Kaustubh Pandit, Fabricio Silveyra, Dan Duong, Clint Hostetler, Ashley Milam, Bradon Nave, Ron Squires, Zhongxin Yu, Kar-Ming Fung, Narendra Battula, Steven Potter, Chongle Pan, Wei R. Chen, Paulo Martins, Yuye Ling, Qinggong Tang

## Abstract

Kidney transplantation remains the preferred treatment for patients with end-stage kidney disease. However, the ongoing shortage of donor organs continues to limit the availability of transplant treatments. Existing evaluation methods, such as the kidney donor profile index (KDPI) and pre-transplant donor biopsy (PTDB), have various limitations, including low discriminative power, invasiveness, and sampling errors, which reduce their effectiveness in organ quality assessment and contribute to the risk of unnecessary organ discard. In this study, we explored the dynamic optical coherence tomography (DOCT) as a real-time, non-invasive approach to monitor the viability of ex vivo mouse kidneys during static cold storage over 48 hours. The dynamic metrics logarithmic intensity variance (LIV), early OCT correlation decay speed (OCDS*_e_*), and late OCT correlation decay speed (OCDS*_l_*) were extracted from OCT signal fluctuations to quantify temporal and spatial tissue activity and deterioration. Our results demonstrate that DOCT provides complementary functional information to morphological assessment offered by conventional OCT imaging, showing potential to improve pre-transplant organ evaluation and clinic decision-making.

## 1. Introduction

For patients with end-stage kidney disease, kidney transplantation is widely recognized as the preferred treatment offering long-term benefits and improved quality of life compared with hemodialysis [1]. After surgical removal, donor kidneys are either preserved in cold storage solution or maintained by a hypothermic perfusion machine, to slow metabolic activity and mitigate the organ deterioration during transportation to the recipient [2]. Once received, the recipient undergoes transplantation surgery followed by careful postoperative renal function monitoring and long-term follow-up care to ensure graft health and overall well-being [3]. Due to multifaceted advances in surgical interventions, immunosuppression regimens, and postoperative management [4], survival rates for kidney transplant recipients have improved substantially [5].

However, a shortage of donor organs continues to limit access to this treatment. As of July 2025, a total of 99,357 patients were on the kidney transplant waiting list in the United States. For reference, a total of 27,759 kidney transplants were performed in the United States in 2024 (Organ Procurement and Transplantation Network, 2025). The gap between limited organ availability and substantial demand continues to place significant pressure, leading to low quantity of life [6], increased mortality [7], and escalating healthcare costs [8]. To alleviate this persistent shortage, expanded criteria donor (ECD) kidneys [9], defined by the United Network for Organ Sharing (UNOS) [10] as organs come from older donors (over 60 years of age) or donors with certain comorbidities, have been included to expand the available donor pool and enable more patients to receive transplants. While efforts have been made, in 2023, kidneys recovered from donors aged 65 years or older had a nonuse rate of approximately 72% [11].

In current clinic practice, the kidney donor profile index (KDPI) serves as an established metric to evaluate deceased donor kidney quality before transplantation [12, 13]. It combines multiple donor factors, such as age, diabetes and hypertension history, serum creatine, height, and weight, to estimate the graft failure risk post transplantation. Although useful for clinical decision making, KDPI has restricted discriminatory capacity [14] as it does not account for individual kidney conditions like kidney biopsy results or cold ischemia time. Consequently, unfavorable frozen-section biopsy results from pre-transplant donor biopsy (PTDB), which is regularly used to evaluate ECD kidneys, becomes the leading reason for organ rejection [15, 16]. PTDB offers valuable information regarding kidney abnormalities [17], such as glomerulosclerosis, necrosis, atherosclerosis, and tubular atrophy, but it carries a risk of post-transplant bleeding [18], and is limited by inferior quality [19-21] compared to standard biopsy preparation, which typically takes several days. In addition, single graft biopsies are prone to sampling errors [22] resulting from evaluating only a small portion of kidney. This poses a risk of unnecessary discard of viable kidneys [23], and has limited predictive power for graft outcomes [24]. In fact, one study reported that if evaluated under the French system, nearly 62% of the kidneys discarded in the United States could have been accepted for transplantation [13]. This underscores the need for complementary tools that provide more reliable evaluation of kidney quality, supporting optimized transplant decisions and outcomes.

A variety of imaging modalities have been explored for ex vivo kidney assessment. Magnetic resonance imaging (MRI) offers volumetric structural and functional information with excellent soft tissue contrast [25], but suffers from high cost and limited accessibility. Hyperspectral imaging is another non-invasive and label-free modality [26]. However, it is limited by shallow penetration depth and high complexity of operation and data processing. Photoacoustic imaging has also been investigated in kidney evaluation by quantifying fibrosis [27], with ongoing efforts to further explore it potential. Optical coherence tomography (OCT) [28] is an non-invasive and staining-free optical imaging modality which utilizes back-scattered light to generate micron-level resolution, cross-sectional tissue images with millimeter-depth penetration. These capabilities make OCT a promising tool for transplant medicine to visualize renal structures [29-32] such as tubules and glomeruli through scanning multiple regions unlike PTDB. However, conventional OCT captures only static anatomical details and cannot reveal metabolic activity, a vital indicator of organ viability and quality [33]. To overcome this, dynamic OCT (DOCT) [34-38] extends conventional OCT by detecting temporal fluctuations in backscattered light driven by underlying cellular motion and metabolic processes. Instead of a single cross-sectional B scan, DOCT acquires repeated scans over time and use motion extraction algorithms [35] to reveal tissue dynamics, which are typically displayed as colormaps. In recent studies, DOCT has been applied to analyze live cell activities across various samples, from human biopsy specimens [39] to mouse tissues such as tongue [40], liver [41], and trachea [36], as well as tumor spheroids [42].

In this study, we focus on investigating the feasibility of using DOCT to monitor tissue dynamics in mouse kidneys at low temperature, during which cellular metabolism is suppressed but remains active [43, 44]. We captured DOCT data from ex vivo mouse kidneys after flushing with preservation solution and storing them at 4°C to simulate the static cold storage [45], and analyzed dynamic signal variations over 48 hours to reveal tissue dynamic activity changes. Given the limitations of current evaluation approaches, this work seeks to provide a reference for future studies utilizing DOCT for pretransplant organ viability and quality assessment, complementing conventional OCT imaging.

## 2. Materials and Methods

### 2.1 Animal Preparation and Kidney Extraction

C57BL/6 mice were purchased from Jackson Laboratories and housed according to the institutional guidelines at the University of Oklahoma (OU). All animal studies were conducted in accordance with the OU Institutional Animal Care and Use Committee (IACUC) regulations. Three mice, aged 4 months, were used in DOCT data acquisition. Following sacrifice, cardiac perfusion was immediately performed using 50–60 mL of University of Wisconsin (UW) preservation solution [46] cooled to 4°C, delivered through the left ventricle by a Smiths Medical Medfusion 3500 syringe pump (see Figure S1a in Supplement 1) at a controlled flow rate to flush out blood. Perfusion continued until the effluent was clear and the kidney appeared visibly pale (Figure S1b). After perfusion, both kidneys were promptly harvested and placed in UW solution at 4°C in a refrigerator (*n* = 6).

### 2.2 OCT System Configuration and Data Acquisition

The OCT imaging system (TEL221PSC1, Thorlabs Inc., Newton, NJ, USA) used in this study provides an imaging depth of 3.5 mm in air (2.6 mm in water), with a lateral resolution of 13 µm and an axial resolution of 5.5 µm in air (4.2 µm in water). A-scan rates range from 5.5 kHz to 76 kHz, with a max sensitivity of 109 dB at 5.5 kHz. The field of view (FOV) was configured to be 4 mm × 1.5 mm in the lateral (*x*) and axial (*y*) scanning directions, with corresponding pixel sizes of 5 µm and 2.54 µm. The A-scan rate was set to 76 kHz, resulting in a B-scan rate of 66.7 Hz.

The mouse kidney was placed in a metal container filled with UW solution during imaging (Figure 1a2) and the container was positioned in a water-filled tray with an ice pack to maintain a low temperature. To capture tissue dynamics across three regions of the kidney, three measurement lines spaced ∼1 mm apart were imaged on each side (Figure 1a3, a4). For each line, 128 sequential temporal B-scan frames were acquired, yielding a total of six measurement lines per kidney. The imaging was performed at 0, 1, 2, 4, 6, 10, 15, 24, 36, and 48 hours post flushing.

**Figure 1.**
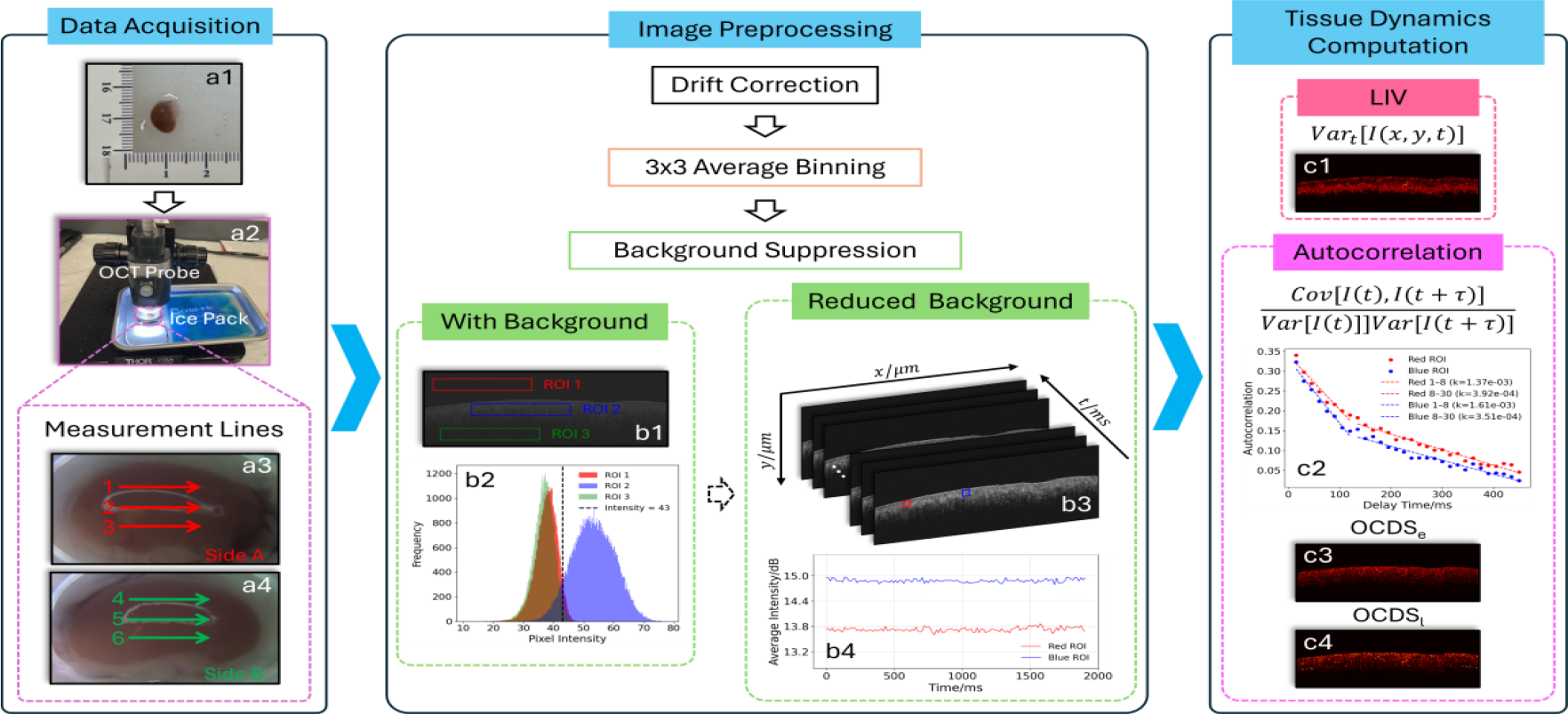
Schematic illustration of the workflow, encompassing data acquisition, image preprocessing, and tissue dynamics computation. (a) DOCT data acquisition: (a1) a kidney from a mouse after flushing; (a2) image capturing; (a3) repeated B-scans along six measurement lines spaced ∼1 mm apart, three on each side of the kidney, to sample different regions. (b) Image preprocessing included drift correction, followed by average binning and background suppression: (b1) 3 ROIs were selected from one OCT intensity frame; (b2) intensity histograms of the three ROIs in (b1) suggest that 43 is an effective threshold for background suppression; (b3) sequential frames from a background reduced DOCT dataset, with two ROIs selected and their average intensities presented in (b4). (c) Tissue dynamics computation: (c1) LIV; (c2) autocorrelation analysis of the intensity values from the two ROIs in (b3), with linear regression slopes computed over the intervals [15, 120] ms and [120, 450] ms, respectively; (c3) OCDS*_e_*; (c4) OCDS*_l_*.

### 2.3 Image Preprocessing Pipeline

To maintain alignment of tissue structures across sequential frames, B-scan drift was first corrected using a cross correction-based method implemented in Fiji [47] via the plugin Fast4Dreg, which incorporates the NanoJ-Core drift correction algorithm [48]. This approach estimates frame-to-frame displacement in the lateral directions (*x, y*) by computing the displacement that best aligns images based on cross-correlation between corresponding maximum intensity projections. The central frame in the sequence was selected as the reference, and all other frames were registered by estimating their translational offset relative to this frame. Data showing uncorrectable drift along the z-axis (orthogonal to the *x*–*y* plane) were excluded. Following drift correction, 3 × 3 average binning without overlap was applied to each frame to reduce noise, mitigate residual artifacts, and improve computational efficiency for subsequent analysis.

A threshold value of 43 was used to separate signal from background, as indicated by intensity histograms derived from three regions of interest (ROIs) in a DOCT intensity frame (Figure 1b1 and 1b2). This threshold was applied to all frames (e.g., Figure 1b3) for subsequent tissue dynamics computation.

### 2.4 Tissue Dynamics Computation

For quantifying the tissue dynamics, the logarithmic intensity variance (LIV) [35] was computed for each pixel to distinguish rapid fluctuations in signal intensity, reflecting localized tissue dynamic activity (Figure 1c1):

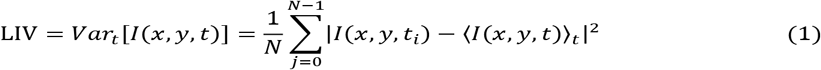

where *N* is the total number of frames in the DOCT dataset (*N* = 128), and *I*(*x, y, t*)>*t* denotes mean intensity value over time *t* for the pixel located at (*x, y*). This method can highlight active regions within tissue while suppressing purely structural signals, thereby effectively demonstrating motion contrast.

However, it lacks the ability to quantify the rate or speed of tissue dynamics, restricting its capacity to differentiate between different dynamic regimes.

To address this limitation, we analyzed two OCT correlation decay speed (OCDS) [35] metrics based on the autocorrelation of the OCT intensity signals:

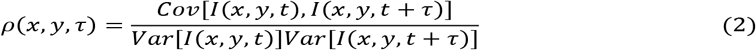

where *Cov* represents the covariance calculation, *Var* denotes the variance, and τ is delay time. Figure 1c2 presents the autocorrelation curves calculated from two ROIs indicated in Figure 1b3. Each point is separated by a time lag of Δτ=15 ms, corresponding to the frame acquisition interval. The first points at τ = 0 was excluded from the analysis. As expected in dynamic samples, the autocorrelation values gradually decrease toward zero with increasing delay time. The OCDS*_e_*, early OCDS, is calculated as the slope of the autocorrelation curve within the 15 ms to 120 ms range to quantify fast tissue dynamics (Figure 1c3). The OCDS*_l_*, charactering slow dynamic behavior, is calculated as the slope of the autocorrelation curve within 120 ms to 450 ms range (Figure 1c4).

### 2.5 Fluorescence Microscopy Imaging

Mouse kidneys were collected following perfusion via the heart to remove residual blood, then preserved at 4°C in UW solution. Imaging was performed under five different conditions: at 0, 6, 24, and 48 hours post flushing, along with an unstained control sample treated with 75% ethanol for 5 minutes to induce cell death, followed by three PBS rinses. Each kidney was cut into small slices (8-10 μm) to facilitate dye penetration. The tissue sections were stained using LIVE/DEAD Viability/Cytotoxicity Kit (Thermo Fisher Scientific, Waltham, MA, USA), in which samples were incubated in PBS containing calcein-AM and ethidium homodimer-1 for 60 minutes at 37°C in the dark, followed by three thorough washes with PBS. Finally, the tissues were mounted on glass microscope slides with PBS, covered with coverslips, and promptly imaged using a BZ-X810 fluorescence microscope (Keyence, Osaka, JAPAN) equipped with 4×/0.10 NA and 20×/0.45 NA objectives. For the 4× objective, the exposure time was 1/30s for the green channel and 1/200s for the red channel, while for the 20× objective, exposure times were 1/150s and 1/1300s, respectively.

### 2.6 Assessment of Regional Heterogeneity

To assess spatial heterogeneity in tissue dynamics across regions, a linear mixed-effects model [49] was fitted at each time point using values from different measurement lines across all mice. A likelihood ratio test [50] was conducted to determine whether there were statistically significant differences among regions, indicating regional variation in tissue dynamic activities.

## 3. Results

Figure 2 presents representative structural intensity images and DOCT colormaps — LIV, OCDS*_e_*, and OCDS*_l_*— of six kidneys acquired one hour after flushing with UW solution. The intensity images reveal overall tissue morphology and exhibit regional variations, reflecting differences in tissue structure and optical scattering properties. The LIV maps highlight localized regions of dynamic intracellular activities, though overall variance remains low at this early post-flush stage. The OCDS*_e_*, which quantifies fast dynamic behavior, shows elevated values in specific regions and highlights localized areas of rapid tissue activities. In contrast, OCDS*_l_*, designed to be sensitive to slower dynamic processes, reveal distinct spatial patterns associated with slower intracellular motions. These differences illustrate the complementary responses of OCDS*_e_*and OCDS*_l_*in concurrently characterizing fast and slow dynamic processes within tissue.

**Figure 2.**
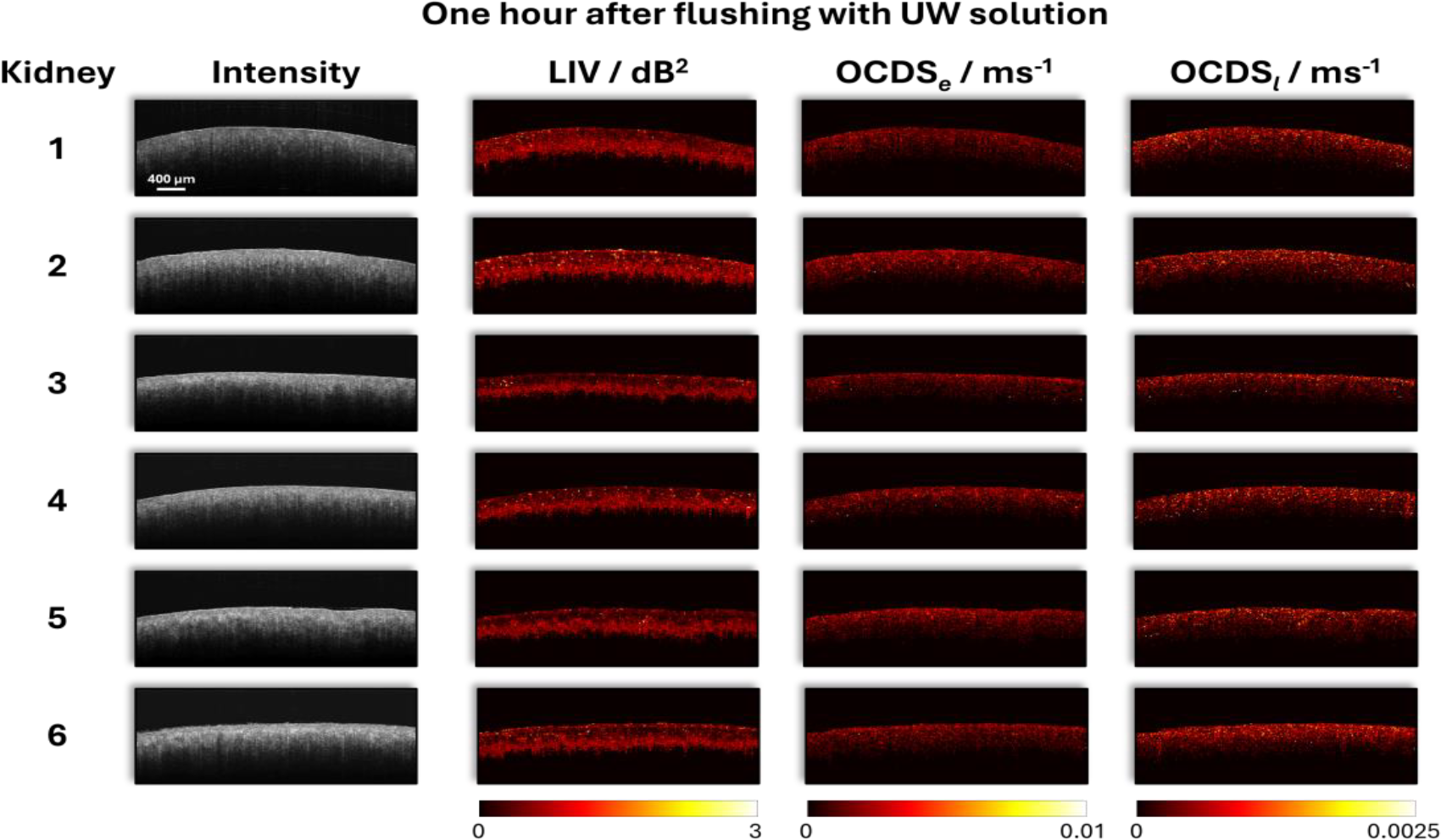
Representative OCT intensity images and corresponding LIV, OCDS*_e_*, and OCDS*_l_*colormaps for six mouse kidneys one hour after flushing with UW solution.

Figure 3 displays the progression of tissue dynamics from 0 to 48 hours after flushing with UW solution. To focus on tissue areas, mean LIV, OCDS*_e_*, and OCDS*_l_*values for each measurement line were computed using pixels with high OCT structural intensity, thereby excluding background and low signal regions. The average and standard deviation across all measurement lines were then calculated for each kidney at each time point to characterize the overall trends and inter-sample variability, as shown in Figure 3a. Generally, LIV values increased rapidly between the 0-hour and 6-hour time points, reaching a maximum before undergoing a sharp decline by the 12-hour time point. Most kidneys then entered a relatively stable phase up to the 36-hour, with a noticeable decrease observed to the 48-hour time point. OCDS*_e_*values showed a general increasing trend, with a quick rise between the 0-hour and 6-hour time points, followed by a slower increase with fluctuations from 6 to 48 hours. Conversely, OCDS*_l_*exhibited a more complex pattern, with fluctuations at first 12 hours followed by a gradual decline though the 48-hour time point. The ridgeline plot in Figure 3b and 3c were generated by aggregating the mean LIV and OCDS*_e_*values, respectively, from each measurement line across all kidneys at each time point. The overall position of each distribution shifts over time, reflecting temporal changes consistent with the trends of mean LIV and OCDS*_e_*curves in Figure 3a.

**Figure 3.**
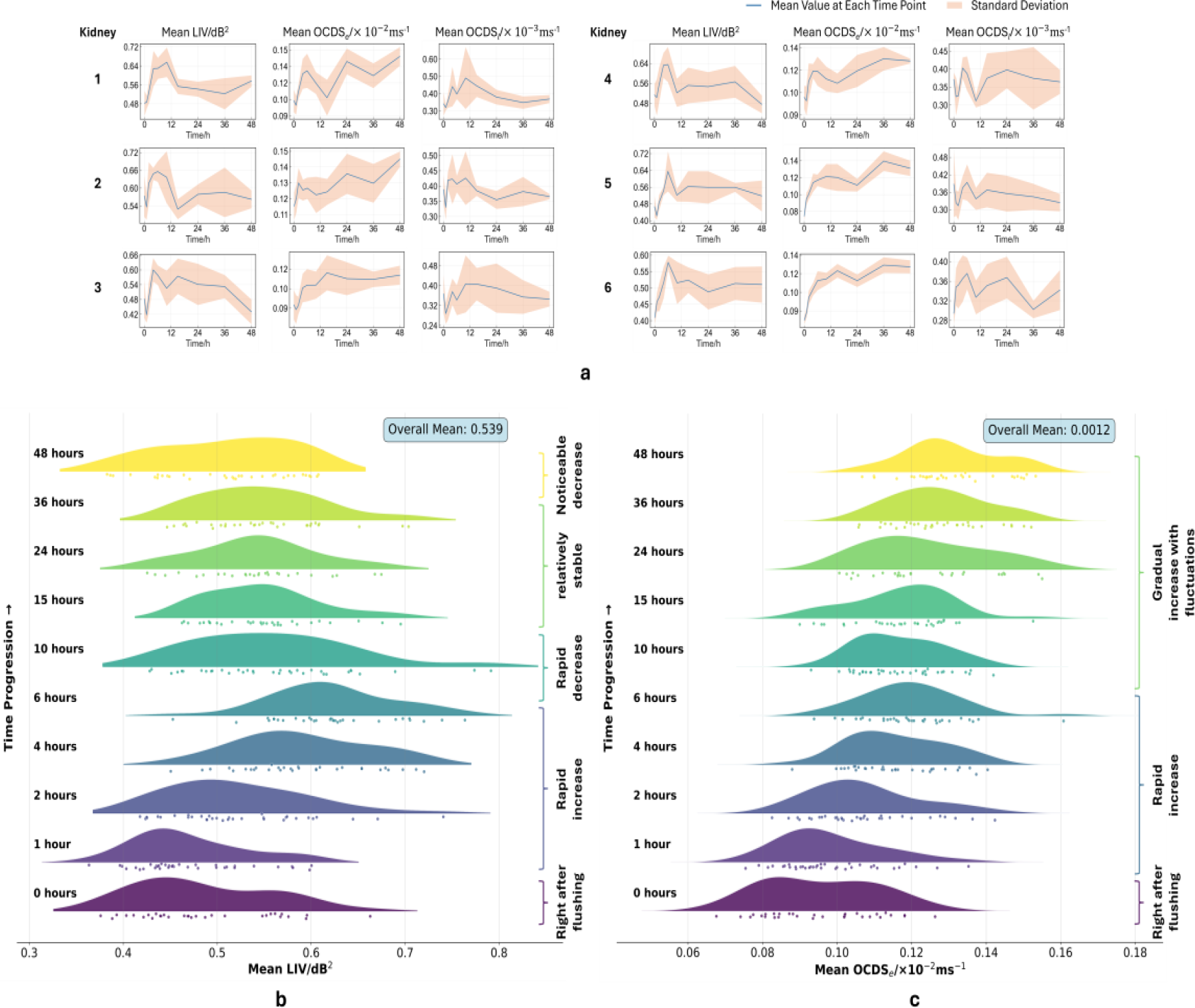
Temporal changes in tissue dynamics within 48 hours after flushing. (a) Mean LIV, OCDS*_e_*, and OCDS*_l_*values for each mouse kidney. (b) Ridgeline plot showing changes in the distribution of mean LIV values across all mouse kidneys over the 48-hour period. (c) Corresponding ridgeline plot for mean OCDS*_e_*values.

To evaluate regional heterogeneity in tissue dynamics, mean LIV values were compared across 6 measurement lines on the kidney at each time point. Figure 4a illustrates an example of LIV colormaps from the second kidney at the 6-hour time point, along with corresponding histograms fitted with lognormal distributions after excluding the lowest and highest 5% of pixel values to reduce the impact of outliers. While the colormaps display distinct spatial patterns, the histograms indicate the distribution of LIV values is consistently well-characterized by a lognormal model. Figure 4b presents a surface visualization of LIV reconstructed from the measurement line data in Figure 4a, illustrating spatial variations across the kidney regions. Figure 4c shows a bar chart of mean LIV values for each measurement line across all kidneys at each time point, and significant differences among measurement lines were observed only at 6-hour time point.

**Figure 4.**
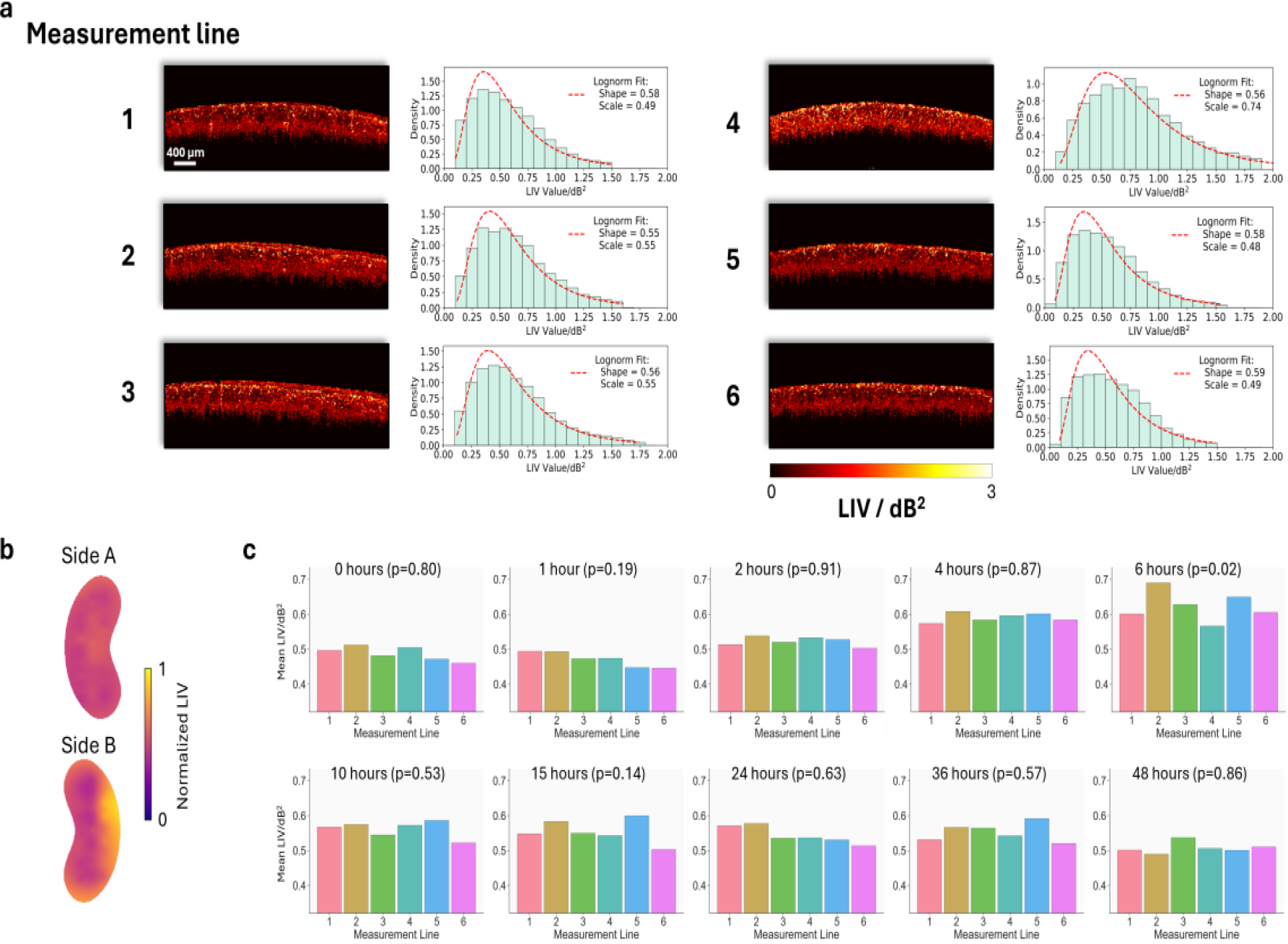
Heterogeneity analysis of LIV. (a) LIV colormaps and corresponding histograms along six measurement lines of the second kidney, acquired 6 hours after flushing. (b) Visualization of LIV distribution on the kidney surface based on data from (a). (c) Mean LIV values for each measurement line across all mouse kidneys at each time point. Significant differences among lines were observed only at the 6-hour time point.

Figure 5 shows fluorescence microscopy images for kidney tissues collected at different time points following UW solution flushing, along with an unstained control group treated with ethanol. Live cells appear green due to calcein-AM staining, a non-fluorescent, cell-permeant dye which is transformed into green-fluorescent celcein by intracellular esterases. The resulting green fluorescence is distributed throughout the cytoplasm across the tissue, producing a diffuse, continuous signal that indicates intact metabolic activity and membrane integrity. In contrast, dead cells appear red due to ethidium homodimer-1, which selectively enters cells with compromised membrane, binds to nucleic acids, and emits red fluorescence. Throughout 0 to 48 hours, all samples displayed green fluorescence, indicating the presence of viable cells and continued metabolic activities. At the same time, dead cells marked by red fluorescence become increasingly dense throughout the observation window. By comparison, the ethanol-treated control group exhibited minimal green intensity and no visible red signal, indicating the presence of intrinsic tissue autofluorescence.

**Figure 5.**
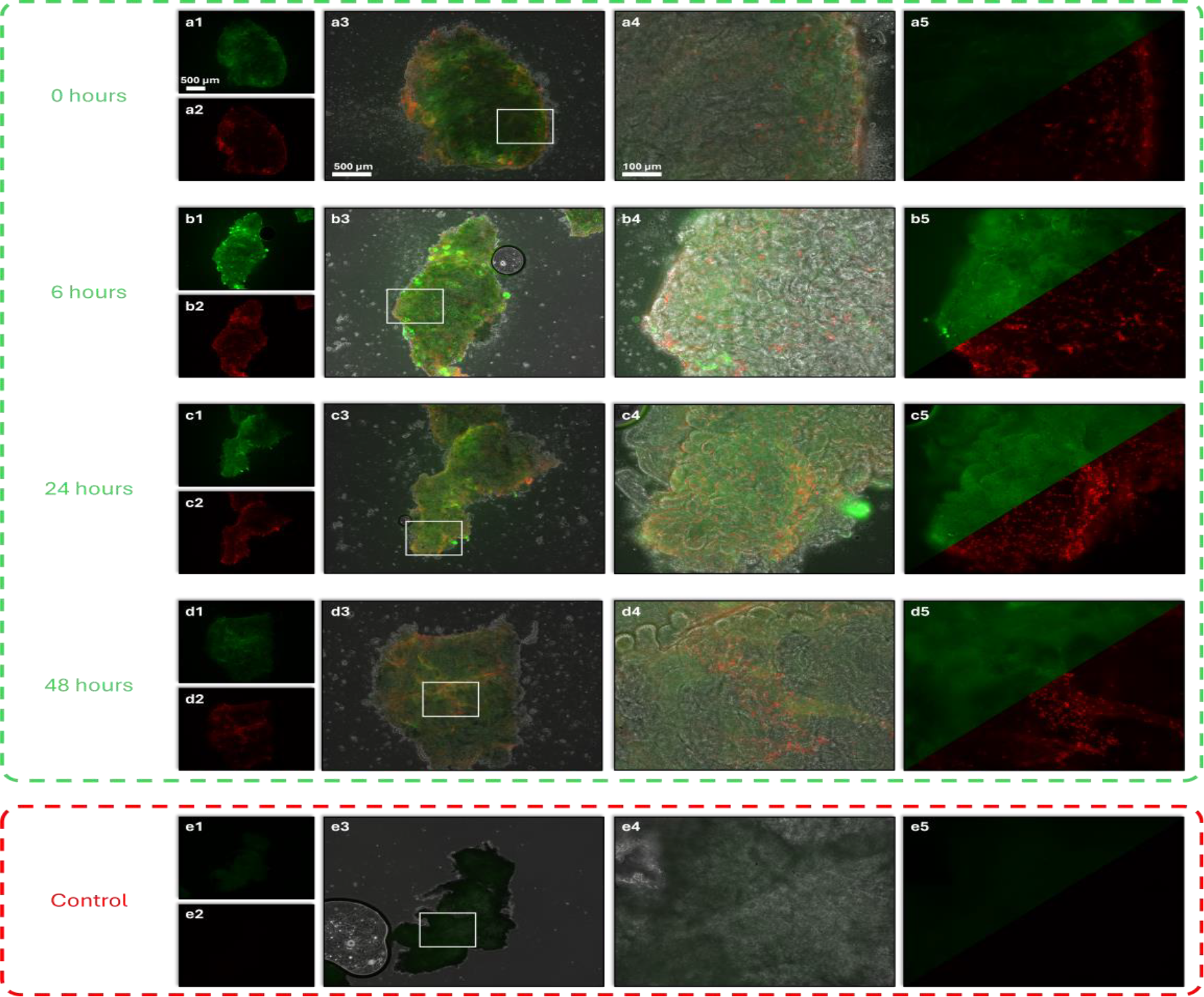
Fluorescence microscopy images highlighting live and dead cells at different time points: (a) 0, (b) 6, (c) 24, and (d) 48 hours. (e) An unstained control group treated with 75% ethanol for 5 minutes to induce cell death. For each condition, there are 5 subpanels: (1) green fluorescence channel indicating live cells at 4× magnification, (2) red fluorescence channel indicating dead cell at 4× magnification, (3) overlayed channels at 4× magnification, (4) overlayed channels of the ROI in (3) at 20× magnification, and (5) 20× magnification with green and red channels occupying the top and bottom, respectively, based on the same field as (4).

## 4. Discussion

In this study, we utilized DOCT to monitor mouse kidney tissue status over a 48-hour period following UW solution flushing. This approach enables noninvasive, label-free assessment of both tissue morphology and dynamic activities at micron-level spatial resolution, providing integrated structural and functional information that conventional methods often fail to provide. By analyzing intensity signal fluctuation patterns to extract quantitative descriptors such as LIV, OCDS*_e_*, and OCDS*_l_*, we evaluated temporal and spatial variations in tissue activity. The mean LIV, OCDS*_e_*, and OCDS*_l_*values displayed distinct trend over the observation period, each reflecting unique aspects of tissue dynamics associated with progressive organ deterioration. Taken together, these metrics provided complementary insights into overall, fast, and slow intracellular motions, offering a functional framework for evaluating tissue viability and degeneration during static cold storage. More importantly, the unique functional information provided by DOCT prior to transplantation may support clinical decision making and improve outcomes. However, several key points require further discussion, as outlined below.

First, from 0 to 6 hours, both mean LIV and OCDS*_e_*showed an increasing trend (Figure 3). This rise may be associated with cellular activity partially recovering from various disturbances [51-53] induced by the left ventricular flush. During this early period, cells benefited from high concentrations of protective (e.g., lactobionic acid and raffinose pentahydrate) and metabolic agents (e.g., adenosine, glutathione, and allopurinol) provided by UW solution, leading to stabilizing processes outweighing necrotic degeneration. This is also consistent with fluorescence microscopy results: at the 0-hour time point (Figure 5a), cells observed mostly inactive with weak green fluorescence due to limited intracellular esterase activity, while only a small number of dead cells were indicated by red fluorescence; whereas by 6 hours, a brighter green fluorescence signal appeared, indicating partially restored cellular viabilities, though with an increasing number of dead cells (Figure 5b). Before the 12-hour time point, a rapid decrease in mean LIV was observed while mean OCDS*_e_*remained at a relatively high level despite some fluctuations. This combination of low LIV and persistently high OCDS*_e_*suggests that necrotic processes were increasing during this phase [35], contributing to the observed changes in DOCT signals. Between 12 and 36 hours, LIV remained generally stable, while mean OCDS*_e_*showed a gradual increase, indicating a mild acceleration of cell death accompanied by relatively steady cellular activities. Beyond 36 hours, mean LIV showed a rapid decline, corresponding to intensified cell death and reduced cellular functions, as indicated in Figure 5d by the dimming of the green fluorescence.

Second, the LIV values were well fitted by log-normal distributions, as shown in Figure 4a. This suggests that the optical fluctuations measured by LIV are from underlying multiplicative tissue activities rather than additive random noise [54], which supports the biological relevance of our DOCT findings. In biological tissue, metabolic activities are determined by multiple interconnected factors rather than by isolated activities, in which those factors are organized in cascaded pathways [55, 56]. As a result, the changes at each step can propagate and be scaled by downstream processes, finally combining to alter tissue optical scattering properties in a way that generates a log-normal distribution observed in DOCT measurements.

Third, the mean LIV values measured along six lines corresponding to six regions on the kidney surface, as shown in Figure 4c, did not show significant differences except at the 6-hour time point, when the most active cellular dynamics were observed (reflected by the highest mean LIV values in Figure 3). This pattern indicates that globally increased cellular activity at 6-hour is not occurring uniformly but rather leading to a high regional heterogeneity. In contrast, the relatively limited heterogeneity at other time points may be attributed to the young age of the mice, which results in less intrinsic variability, and to the measurement lines spanning large areas on the kidney surface, which likely averages out local fluctuations in cellular dynamics.

Finally, cells remained viable at the 48-hour time point, as indicated by the presence of measurable LIV, OCDS*_e_*, and OCDS*_l_*signals (Figure 3) and confirmed by the live/dead viability test (Figure 5). In addition, as shown in Figure S2 in Supplement 1, the mean values of these dynamic signals measured in formalin-fixed mouse kidneys were noticeably lower than those observed at 48 hours. This further supports the dynamic optical signals captured by DOCT contain information reflecting cellular metabolism.

In summary, our findings demonstrate DOCT, including LIV, OCDS*_e_*, and OCDS*_l_*, can capture tissue dynamics related to cellular metabolism, reflecting organ status in real time. These dynamic signatures can complement the structural information provided by conventional OCT imaging, which has already been investigated for predicting post transplantation outcomes and supporting clinical decision making and surgical management. This approach offers new avenues for organ quality evaluation prior to transplantation and noninvasive assessment during organ preservation.

Although this study provides valuable insight into application of DOCT on organ tissue dynamic activity monitoring, there are several limitations. First, the sample size is limited, which may undermine the generalizability of the findings. Second, all measurements were conducted on kidneys from young mice under controlled ex vivo conditions, which limited the ability of this study to represent the more complex scenarios encountered in human organs. Third, while LIV, OCDS*_e_*, and OCDS*_l_*characterize tissue dynamics, there are also fast Fourier transform (FFT) based methods that can separate the dynamic signals into different motion components, such as slow, medium, and fast, enabling a more comprehensive analysis and detailed characterization of tissue viability. Despite the limitations, this study demonstrates the feasibility of using DOCT to capture dynamic tissue activity to assess the organ viability.

## Acknowledgement

This work was supported by grants from the University of Oklahoma Health Sciences Center (P30CA225520), National Science Foundation (OIA-2132161, 2238648, 2331409), National Institute of Health (R01DK133717), Oklahoma Center for the Advancement of Science and Technology (HR23-071), the medical imaging COBRE (P20 GM135009), the Prevent Cancer Foundation, a grant from the Data Institute for Societal Challenges and the Research Council funded by the Office of the Vice President for Research and Partnerships of the University of Oklahoma Norman Campus, and the Midwest Biomedical Accelerator Consortium (MBArC), an NIH Research Evaluation and Commercialization Hub (REACH). Financial support was provided by the OU Libraries’ Open Access Fund.

## Disclosures

The authors declare no competing interests.

## Data Availability

The data that support the findings of this study are available from the corresponding author upon reasonable request.

## Notes

### Competing Interest Statement

The authors have declared no competing interest.

