## Supplementary Figures for "Real-Time Viability Assessment of Ex Vivo Mouse Kidneys for Transplant Applications Using Dynamic Optical Coherence Tomography"

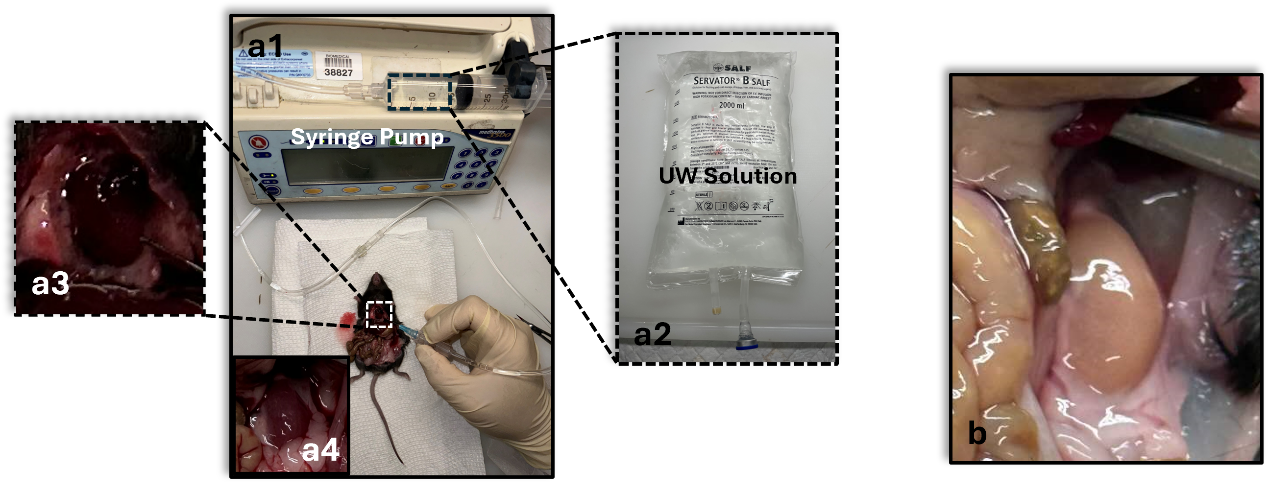

Figure S1. Experimental setup used for kidney flushing with UW solution. (a1) Experimental setup. (a2) UW solution. (a3) Insert the needle into the left ventricle. (a4) A kidney before flushing. (b) A kidney after flushing.

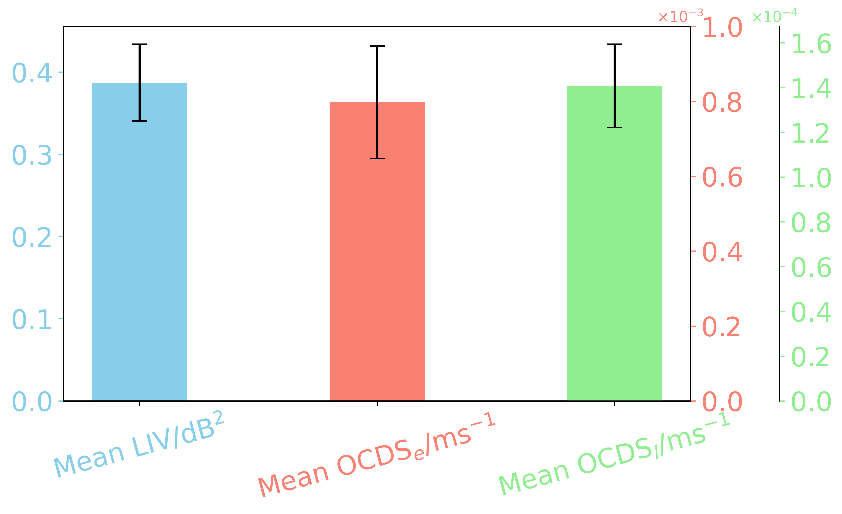

Figure S2. Mean LIV, OCDS*_e_* and OCDS*_l_* values of formalin-preserved mouse kidneys The overall mean values across all mice were 0.387, 7.96×10⁻⁴, and 1.41×10⁻⁴, respectively.
